# Proteomic signatures of acute oxidative stress response to paraquat in the mouse heart

**DOI:** 10.1101/2020.05.29.124487

**Authors:** Vishantie Dostal, Silas D. Wood, Cody T. Thomas, Yu Han, Edward Lau, Maggie P.Y. Lam

## Abstract

The heart is sensitive to oxidative damage but a global view on how the cardiac proteome responds to oxidative stressors has yet to fully emerge. Using quantitative tandem mass spectrometry, we assessed the effects of acute exposure of the oxidative stress inducer paraquat on protein expression in mouse hearts. We observed widespread protein expression changes in the paraquat-exposed heart especially in organelle-containing subcellular fractions. During cardiac response to acute oxidative stress, proteome changes are consistent with a rapid reduction of mitochondrial metabolism, coupled with activation of multiple antioxidant proteins, reduction of protein synthesis and remediation of proteostasis. In addition to differential expression, we saw evidence of spatial reorganizations of the cardiac proteome including the translocation of hexokinase 2 to more soluble fractions. Treatment with the antioxidants Tempol and MitoTEMPO reversed many proteomic signatures of paraquat but this reversal was incomplete. We also identified a number of proteins with unknown function in the heart to be triggered by paraquat, suggesting they may have functions in oxidative stress response. Surprisingly, protein expression changes in the heart correlate poorly with those in the lung, consistent with differential sensitivity or stress response in these two organs. The results provide insights into oxidative stress responses in the heart and may avail the search for new therapeutic targets.

## Background

Oxidative stress is widely implicated in cardiovascular disorders and other age-related diseases [1,2]. Evidence of oxidative stress is commonly found in many animal models of heart diseases, but antioxidant therapeutics have only seen limited success and were in some cases harmful, suggesting further work is needed to understand stress response pathways and identify new therapeutic targets [3,4]. It is now known that acute oxidative stress triggers complex and multifactorial changes to the cardiac proteome including widespread chemical and enzymatic oxidative modifications [5,6] as well as the activation of integrated stress response pathways [7]. Prior proteomics data on the global responses of cultured mammalian cells to hydrogen peroxide in vitro pointed to close connections between oxidative stress and the unfolded protein response, as well as a diverse number of regulatory pathways involved in metabolism and protein translation [8]. Whether similar proteome-wide responses are recapitulated in the heart of in vivo animal models remains unclear.

Despite the biological significance of oxidative stress to heart diseases, to our knowledge there have been few reports on the effect of direct acute oxidant exposure on protein abundance in the heart. To gain insights into how acute oxidative stress remodels the heart proteome, we examined the effect of acute exposure to paraquat as a model of oxidative stress in the mouse. Paraquat (N,N-dimethyl-4,4-bipyridinium dichloride) is a powerful pro-oxidant that generates mitochondrial reactive oxygen species in situ at the respiratory chain [9–11]. Acute paraquat exposure leads to significant multi-organ toxicity and mortality [12] and in laboratory experiments has been widely used as a model of oxidative stress in the heart [13,14]. We therefore compared the cardiac proteome between normal and paraquat-stressed mouse hearts using quantitative mass spectrometry.

## Results

### Moderate to high doses of paraquat induces acute oxidative stress responses in the heart

Paraquat is a mitochondria-targeted redox-cycler that acts as a superoxide generator to produce ROS through interactions with Complex I within the inner mitochondrial matrix [11]. To evaluate the proteomic responses to elevated oxidative stress, we first administered various doses of paraquat to C57BL/6J mice for 24 hours. The animals (n=4 per group) were exposed to three doses of paraquat (low – 10 mg/kg; moderate – 50 mg/kg; and high – 75 mg/kg). The doses were chosen from a range previously reported for cardiac research models in the literature, with reports showing that exposure at 40–80 mg/kg for up to 48 hours induced acute oxidative stress responses as well as reduced contractile function and calcium handling in the mouse heart [13,15–18]. We first compared whether the applied paraquat doses elevated protein abundance of previously documented paraquat induced genes in the heart using mass spectrometry (**Figure 1a**). Pyruvate dehydrogenase kinase 4 (PDK4) is a potent inhibitor of pyruvate conversion into acetyl-CoA that reduces oxygen consumption in the respiratory chain and has previously been found to be strongly induced by 50 mg/kg paraquat [19] as well as the related dipyridyl diquat [20]. Under moderate to high doses but not the low dose of paraquat, we observed a strong induction of PDK4, in addition to the products of two other known paraquat-induced cardiac genes, metallothionein-1 (MT1) and BCL2-like 1 (BCL2L1).

**Figure 1.**
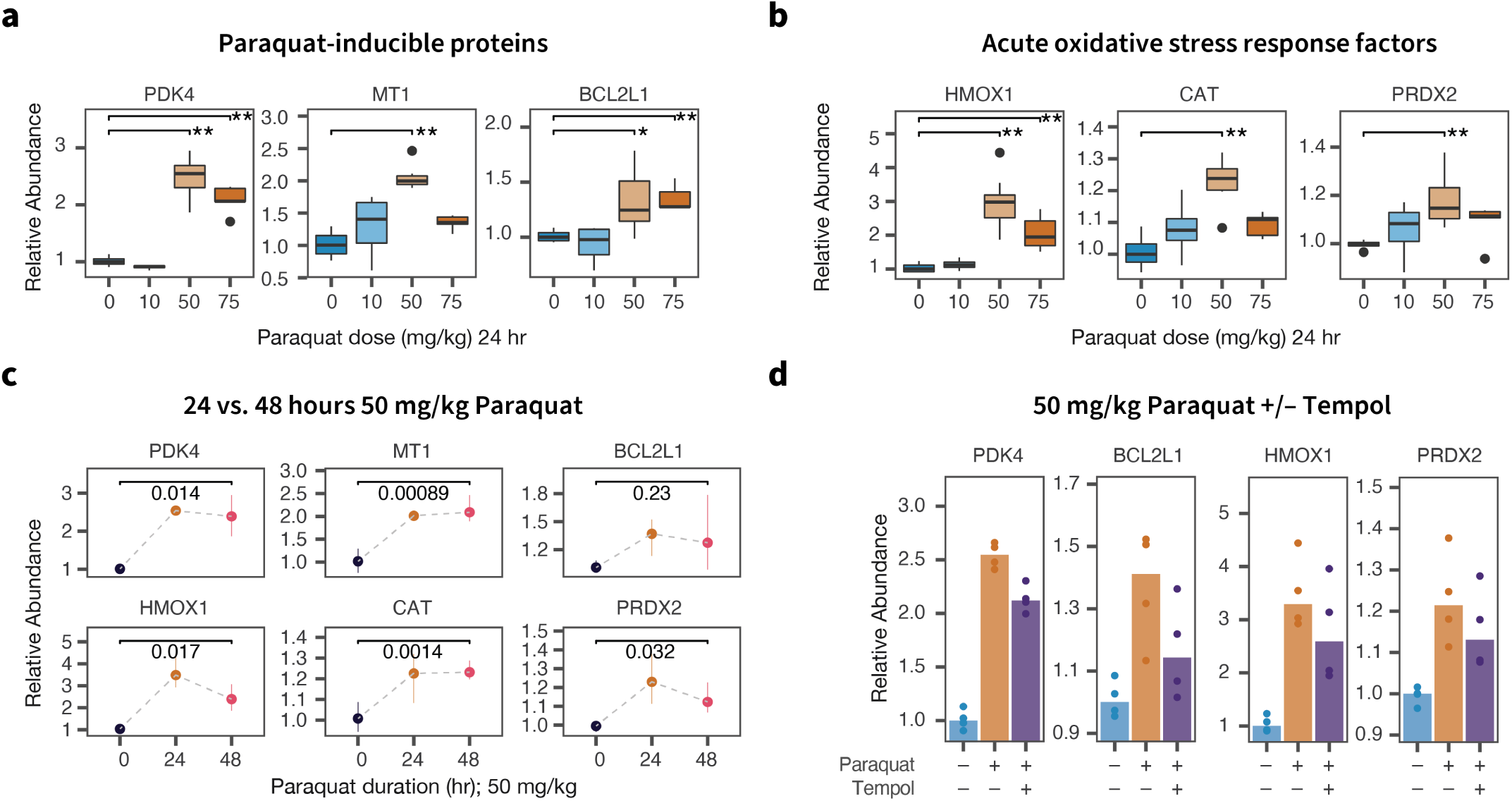
Moderate and high doses of paraquat induce acute oxidative stress response proteins in the heart. **a**. Moderate and high doses of paraquat for 24 hours led to an increase in protein abundance of known paraquat-induced genes in the hearts including PDK4, MT1, and BCL2L1;*: t test P < 0.05; * *: 0.005 vs. vehicle. **b**. Moderate and high doses of paraquat for 24 hours led to a robust increase in heme oxygenase 1 (HMOX1) as well as a moderate increase in the protein level of other NRF2-induced acute oxidative stress response elements in the heart; *: t test P < 0.05; * *: 0.005 vs. vehicle. **c**. A number of induced proteins remained elevated at 48 hours after the paraquat dose including PDK4, MT1, and CAT, whereas other acute response elements including HMOX1 began to subside; numbers: t test P values between 48 hrs vs. vehicle. **d**. Pre-treatment of the anti-oxidant compound Tempol partially repressed the up-regulation of paraquat-induced and acute oxidative stress response proteins.

We next compared whether the paraquat doses employed in the experiment activated oxidative stress responses in the heart as expected. In the heart and in other organs, acute oxidative stress response is under the master regulatory control of the nuclear respiratory factor 1/2 (NRF1/2) pathway [21,22]. Under physiological conditions, NRF2 is continuously degraded and has a short half-life, but under stress conditions NRF2 accumulates and translocates into the nucleus to activate genes with antioxidant response element (ARE) sequences and thereby orchestrates the acute phase of cellular oxidative stress responses. At 24 hr, we observed a robust induction of heme oxygenase 1 (HMOX1), a primary cardioprotective oxidative stress response protein in the heart (**Figure 1b**). At the same time, other NRF2-induced factors were modestly elevated including catalase (CAT), glutathione dismutase 1 (GPX1), and glutamate-cysteine ligase (GCLC/GCLM) (**Figure S1a**). In our experiment, the induction of stress response genes was the highest at 50 mg/kg, hence we selected this dose to further assess the effect of exposure duration and co-administered antioxidant. Contrasting the effects at 24 vs. 48 hours of paraquat exposure, we further found that the assessed paraquat signatures PDK4, MT1, and BCL2L1 remained elevated at 48 hours after paraquat exposure, but several NRF2-induced oxidative response proteins regressed partially toward the baseline, with the exception of catalase, suggesting the acute phase response may begin to subside 48 hours after paraquat administration (**Figure 1c; Figure S1b**). Pre-treatment with Tempol (2 mM; 24 hr) prior to 50 mg/kg paraquat exposure for 24 hours partially reverted the oxidative stress response and paraquat-induced proteins toward the baseline, consistent with the observed protein changes arising from bona fide oxidative stress response induced in the animal models (**Figure 1d; Figure S1c**; also see section on Responses to Antioxidant Treatments).

### Compartment-specific proteome response to acute oxidative stress

The initial results above confirmed that exposure to moderate to high doses (50–75 mg/kg) of paraquat for 24 to 48 hours robustly activated classic acute oxidative stress responses in the mouse heart, and at the same time induced other paraquat-specific signatures. To further investigate how acute stress responses remodel the global cardiac proteome, we selected the medium dose paraquat exposure (50 mg/kg, 24 hr) and acquired deep quantitative proteome profiles using subcellular fractionation and twodimensional peptide separation (**Figure 2a**). To improve coverage, we used a commercial differential lysis workflow to perform subcellular fractionation on the harvested tissue which created three distinct fractions (S1, S2, and S3) containing sets of proteins distinguished by localization and solubility. The samples from each treatment group, tissue, and fraction were randomized and labeled with stable isotopes for quantitative mass spectrometry (**Figure S2a–d**). From the isobaric-labeled quantitative proteomics data, we quantified 5,323 non-redundant protein groups to compare the expression of protein groups in the hearts of vehicle vs. paraquat treated mice. Identified proteins are distributed over diverse cellular compartments (**Figure 2b**). We compared the fraction compositions and found distinct sets of proteins enriched in each fraction (**Figure 2c**). Gene annotations of enriched proteins showed that the S1 fraction is enriched in cytosolic components, whereas both the S2 and S3 fractions are enriched in mitochondria and membrane proteins over the S1 fraction (**Figure S3a**). When comparing the S2 and S3 fractions directly, we further found that the S2 fraction is relatively enriched in myofibrils and nuclear proteins whereas the S3 fraction is enriched in endoplasmic reticulum and sarcolemmal proteins (**Figure S3b**) and overall has more hydrophobic proteins (**Figure S3c**), suggesting the subcellular fractionation was effective in separating the cardiac proteomes into distinct fractions.

**Figure 2.**
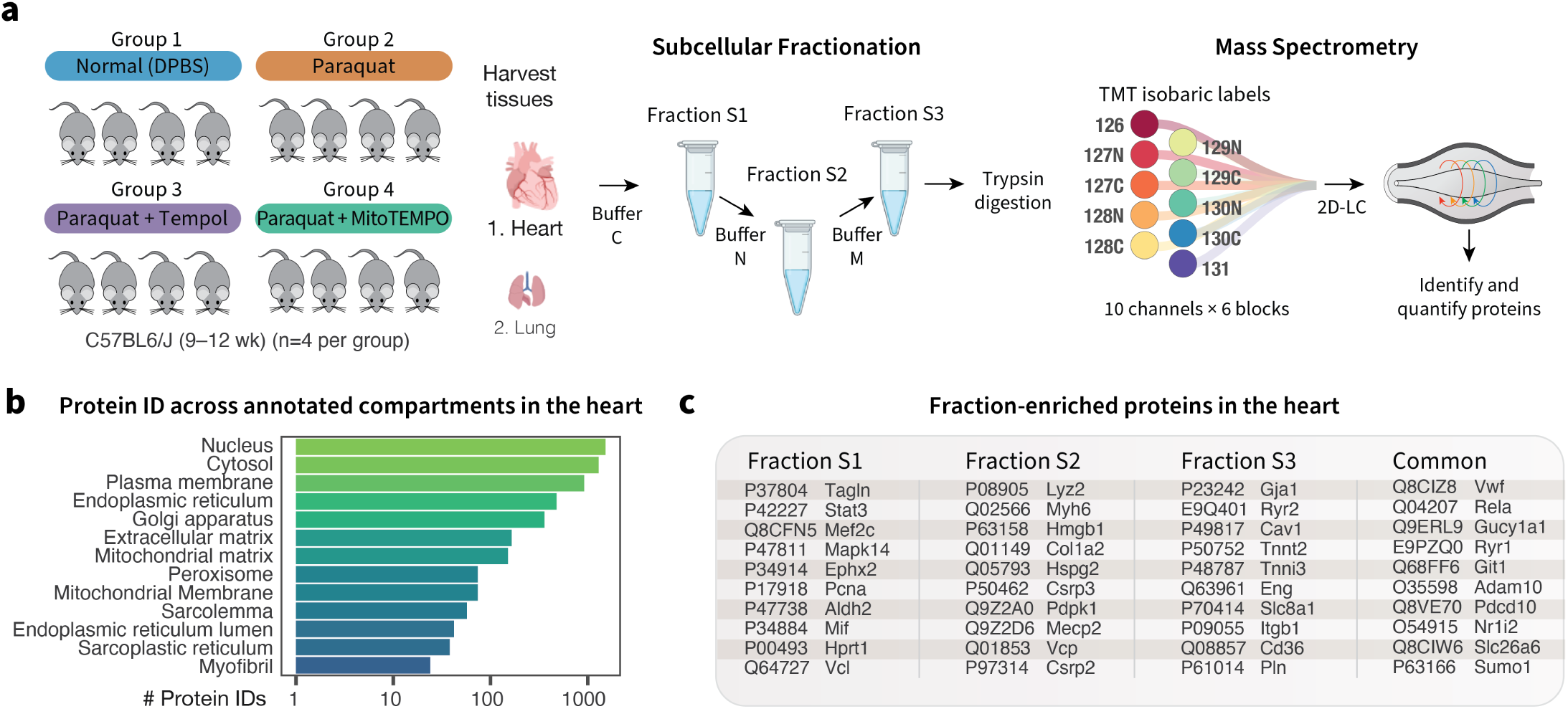
Mass spectrometry analysis of protein expression in normal mouse hearts and following acute paraquat exposure. **a**. Experimental design. C57BL/6J mice (n=4 per group) were treated with a vehicle (DPBS) or a moderate dose (50 mg/kg) of paraquat for 24 hours. Additional groups were given the anti-oxidants Tempol and MitoTEMPO. Cardiac tissues were harvested, homogenized and further separated into three subcellular fractions (S1, S2, S3) using commercially available differential lysis buffers. The resulting proteins were digested and labeled with isotope tags, then combined for protein identification and quantification. **b**. Number of proteins identified in the proteomics experiment across selected cellular compartments. **c**. The subcellular fractions are enriched in different cellular compartments and proteins. Selected proteins relevant to cardiovascular research that are highly enriched in each fraction (*≥* 2-fold, limma adjusted P *≤* 0.01) or are shared across all fractions are shown.

**Figure 3.**
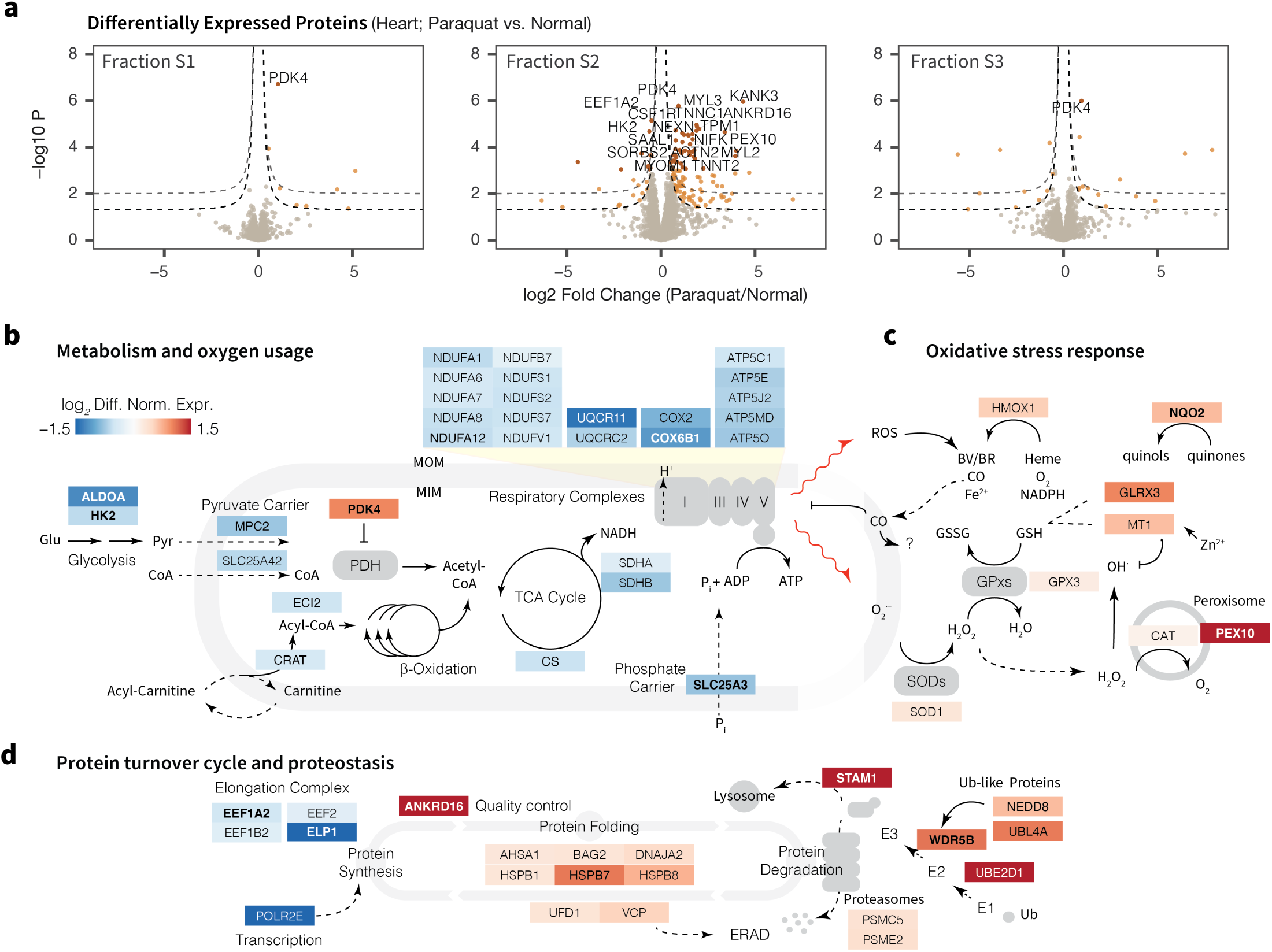
Proteomic signatures of acute oxidative stress in the heart. **a**. Paraquat preferentially causes the differential expression of proteins in the S2 fraction, which is enriched in proteins in the mitochondria, endoplasmic reticulum, and nucleus. Dashed lines: nominal significance at absolute logFC of 0.5 and P ≤ 0.05 and ≤ 0.01, respectively. Brown data points are significantly differentially expressed at 10% false discovery rate; up to 30 differentially expressed proteins are labeled. The pathway diagrams illustrate the differential expression of proteins involved in **b**. central metabolism and respiration, **c**. oxidative damage response, and **d**. protein turnover and proteostasis. Colors represent fold change in paraquat vs. normal hearts, proteins labeled in bold correspond to proteins within the 10% false discovery rate filter in the panels above. The proteomic profiles in stressed hearts are indicative of decreased respiration and protein synthesis coupled with increases in antioxidant proteins and protein degradation amid acute paraquat challenge.

Surprisingly, we found that 50 mg/kg paraquat primarily induced differential expression of proteins localized in Fraction S2, with far fewer differentially regulated proteins in fractions S1 and S3 (**Figure 2a**). Upon closer inspection of the data, we found that acute paraquat exposure broadly induced the differential regulation of a number of mitochondrial and endoplasmic reticulum proteins in the S2 fraction. In particular, we found that the primary feature of the paraquat remodeled proteome appears to be a broad down-regulation of mitochondrial central metabolism proteins, including components of the respiratory chain and the tricarboxylic acid cycle (**Figure 2b**). Concomitant with the downregulation of metabolic proteins is a conspicuous up-regulation of PDK4, which serves to inhibit pyruvate influx into the mitochondrial respiratory chain, suggesting that a principal response of the heart to acute oxidative stress is in the suppression of central metabolism and reduction of respiratory oxygen usage.

At the same time, we observed prominent changes in oxidative stress response proteins (**Figure 2c**). As above, HMOX1 is induced following paraquat exposure. HMOX1/HO-1 is an inducible isoform of heme oxygenase and a prominent NRF2-dependent factor in oxidative stress response both in the heart and in other tissues under general oxidative stress conditions [23]. HMOX1 degrades heme into iron, CO, and biliverdin, the latter being an antioxidant that scavenges peroxyl radicals [24]. MT1 is also upregulated across all examined fractions. Metallothioneins are efficient chelators of zinc and other heavy metal ions and serve as efficient scavengers of hydroxyl and superoxide radicals [25]. Thirdly, we found evidence of a coordinated proteostasis response through a reduction in global protein synthesis and promotion of protein folding and protein degradation (**Figure 2d**). A number of elongation complex proteins including EEF1A2 and EEF2 are down-regulated, whereas protein quality control factors are up-regulated. Among the induced chaperones is HSPB7, a cardiac heat shock protein that is linked to cardiomyopathy in genome-wide association studies and significantly co-occurs with heart failure publications in the literature. However, not every integrated stress response component showed evidence of regulation from the differential expression data. For instance, we did not observe clear trends for differential regulation in mitochondrial import which forms an integral part of the mitochondrial unfolded protein response.

Lastly, the proteomics data also uncovered the differential expression of a number of proteins with still unclear functions in the heart, implicating a potential function of these proteins in acute oxidative stress responses. For example: (i) Proline rich protein 16 (PRR16) is significantly activated after paraquat (logFC: 1.77, P: 1.3e–4; adjusted P: 0.019). Promoter binding analysis provides supporting evidence that the *PRR16* gene may be a target of CREB which is activated by ER stress to trigger unfolded protein response. Although our experiment did not identify CREB1, CREB coactivator CRTC2 is also robustly increased (logFC: 1.06, P: 3.4e–3, adjusted P: 9.9e–2), suggesting PRR16 may be a downstream ER stress response factor. The function of PRR16 in the heart is unclear. (ii) Aldehyde dehydrogenase 16 family member A1 (ALDH16A1) is nominally up-regulated under paraquat exposure (logFC: 1.47, P: 2.0e–2; adjusted P 2.5e–1). Although a member of the aldehyde dehydrogenase family, ALDH16A1 contains aldehyde dehydrogenase domains that are thought to be inactive [26], and its function in the heart remains unknown. It is ubiquitously expressed and has no clear co-expressed genes across human tissues, but previous ChIP-seq data has shown that its promoter region may bind to STAT3 [27], which is also upregulated in the data (logFC: 0.65, P: 3.8e–4; adjusted P: 2.9e–2). (iii) Signal transducing adaptor molecule (SH3 domain and ITAM motif) 1 (STAM) is conspicuously up-regulated following paraquat exposure (logFC 4..00; P: 1.4e–4; adjusted P: 1.9e–2) and is believed be involved in cell growth [28] and vesicular trafficking [29], and is associated with brain neoplasm in the literature. Taken together, the data portray wide-ranging pathway-dependent and compartment-dependent consequences of paraquat exposure on the cardiac proteome, and nominate a number of novel acute response proteins for further characterization.

### Changes in subcellular fraction localization in acute oxidative stress

Myofibrillar disarray and alterations in cytoskeleton dynamics are commonly observed in oxidative damage models. To assess whether we could detect this process or other redistribution of proteins from the proteomics data, we examined whether some proteins can be observed to have differential subcellular fraction distribution in normal vs. paraquat treated hearts. We saw that overall the subcellular fraction distribution between the S2 and S1 fractions, and between the least soluble S3 and S1 fractions, are highly correlated in normal and paraquat samples (Spearman correlation coefficient *ρ* : 0.907 and 0.943), indicating the overall fractionation process is reasonably reproducible (**Figure 4a**). However, there was also a small subset of proteins that appeared to show altered subcellular distribution between the partially overlapping S2 and S3 fractions (*ρ* : 0.521) (**Figure 4b**). We therefore performed a direct comparison of changes in protein ratios between the S2 and S3 fractions in paraquat-stressed hearts over the S2-to-S3 ratio in the normal heart of using the linear model in limma. Among the proteins with significant differences are a number of sarcomeric proteins including myosin and troponin (**Figure 4c**).

**Figure 4.**
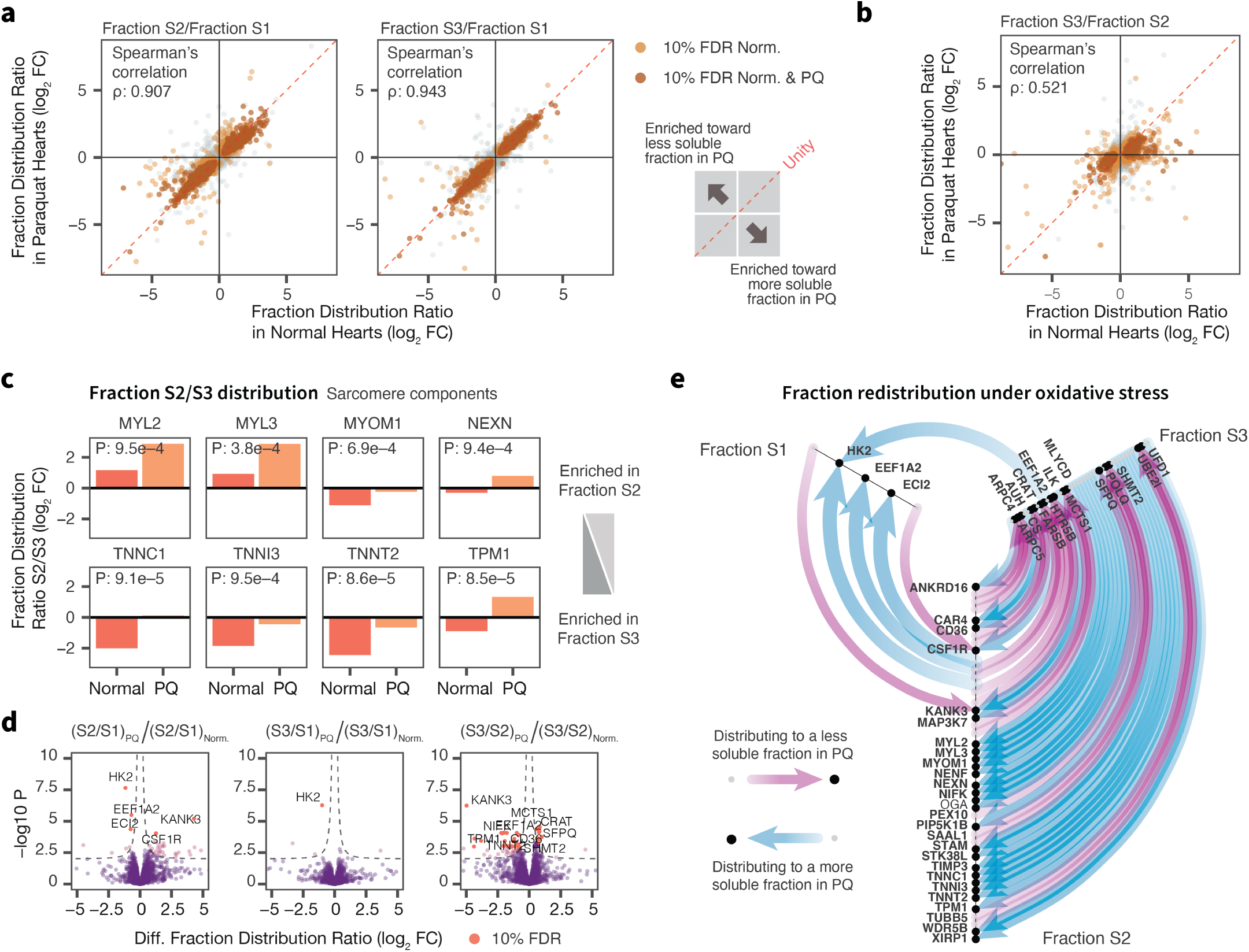
Protein fraction redistribution upon oxidative stress treatment. **a**. The majority of proteins showed consistent fraction localization in normal (x) vs. stressed hearts (y) when comparing the S2 or S3 fraction to the more soluble S1 fraction (Spearman’s correlation *ρ* : 0.907 and 0.943, respectively), but a subset of proteins showed evidence of preferential distribution to more soluble or less soluble fractions upon stress. Each data point is a protein; color: significant differences in protein abundance across fractions in normal (light brown) or both normal and stressed hearts (dark brown). **b**. Correlations in relative protein distribution between the S2 and S3 fractions in normal vs. stressed heart (Spearman’s correlation *ρ* : 0.521). **c**. Bar charts of fraction S2/S3 abundance ratios showing a number of sarcomeric proteins with preferential redistribution from the S3 to the S2 fraction in paraquat (PQ) stressed hearts over normal hearts. Numbers: t test P values. **d**. Volcano plots showing significantly redistributed proteins in stressed hearts (x: log2 FC; y: –log10 P). Red data points are significantly different at 10% FDR in limma; up to 10 of the top significant data points are labeled. **e**. Flow diagram showing redistribution of proteins in stressed hearts among the three subcellular fractions at 10% FDR. Nodes: proteins distributed among the axes representing 3 subcellular fractions; purple edges: redistribution to a less soluble fraction; blue edges: redistribution to a more soluble fraction.

A question is whether we could distinguish translocation from differential expression. A plausible alternative explanation for the increased ratios of S2/S3 for sarcomeric proteins in paraquat treated hearts would be a rapid increase in the expression of sarcomeric genes, coupled with changes in fractionation efficiency; however, given these proteins are abundant and have long half-life in the mouse heart [30], we believe massive changes (e.g., doubling in proteome size) within 24 hours is unlikely and the more parsimonious explanation of the data would be that it is consistent with the dissolution of myofibrils coupled with appearance in the more soluble fraction. Furthermore, we likewise observed a small but significant number of potential translocation events including prominently hexokinase 2 (HK2) (**Figure 4d**). HK2 is known to translocate between the cytosol and mitochondria in the heart following various pathological stimuli [31,32]; thus the result is consistent with at least some of the fraction relocalization proteins representing bona fide spatial translocation. Overall, we observed a complex redistribution of 45 proteins across the subcellular fractions during acute oxidative stress (**Figure 4e**). Although further establishing the cellular localizations of the proteins will require a different experimental design such as using differential ultracentrifugation, our results nevertheless highlight the opportunity to observe potential protein translocation events as a dynamic aspect of the acute phase proteome response to oxidative stress.

### Responses to global and mitochondria-targeted antioxidant treatments

To assess whether antioxidant treatment could rescue the detected proteome perturbations, we compared protein expression between paraquat stressed hearts and antioxidant treatments. Mice in the antioxidant treatment groups were administered with either the cytosolic antioxidant Tempol (4-hydroxy-TEMPO or 4-hydroxy-2,2,6,6-tetramethylpiperidin-1-oxyl) (2 mM in drinking water, ad libitum; 24 hr) [33] or its triphenylphosphonium-conjugated lipophilic variant MitoTEMPO which localizes to the mitochondrion [34] (0.7 mg/kg, i.p.). Considering an arbitrary set of proteins with apparent upward or downward trends following paraquat exposure (P≤ 0.01; | logFC |≥ 0.3), we found that the majority of these proteins reverted toward baseline, including 84% in the Tempol group and 63% in MitoTEMPO group. However, the reversal was mostly partial in magnitude and few reached statistical significance on an individual protein level after multiple-testing correction. Overall, approximately 60% of differentially expressed proteins showed some sign of reversal toward the baseline in both antioxidant groups whereas about 13% of proteins showed no evidence of reversal in our threshold (**Figure 5a**).

**Figure 5.**
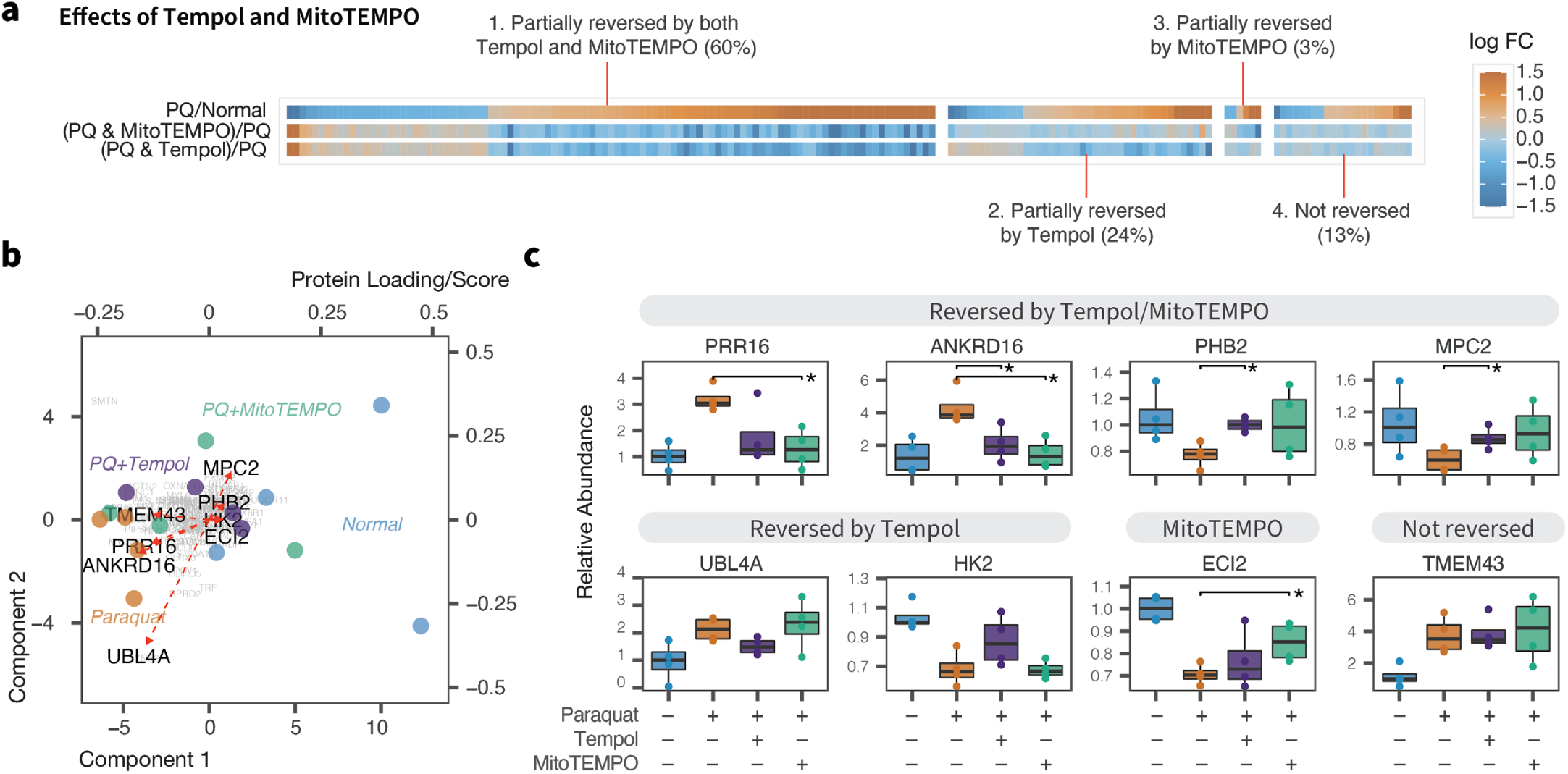
Partial reversal of proteome remodeling by antioxidants. **a**. Heat map showing the fold changes 173 cardiac proteins with apparent changes in abundance following paraquat (PQ) treatment (P < 0.01; | logFC | ≥ 0.3) and their fold changes when comparing co-treatment with either antioxidant Tempol or MitoTEMPO with paraquat to paraquat only. The majority of proteins showed partial reversal to the baseline. **b**. First two components and overlaid protein loadings in linear discriminant analysis between normal, paraquat, and paraquat co-treated with either antioxidants. **c**. Box plots showing relative abundance of selected proteins, in normal, paraquat, paraquat + Tempol, and paraquat + MitoTEMPO hearts. * : t test P < 0.05.

We also observed evidence that some protein changes were preferentially rescued by one antioxidant over the other. To investigate further, we used a linear discriminant analysis to identify proteins with high weights in discriminating between normal, stressed, and antioxidant treated samples (**Figure 5b**). We found that whereas a number of protein changes including PRR16, ANKRD16, PHB2, MPC2 were rescued by both Tempol and MitoTEMPO, UBL4A and HK2 were more reversed in Tempol, and other proteins including TMEM43 were not reversed (**Figure 5c**). Although further investigations will require comparisons of doses, delivery methods, and pharmacokinetics, the data here hint at potential qualitative differences between Tempol and MitoTEMPO, and identify stress-responsive proteins that are not reversed by antioxidant treatments which could be important for the interpretation of antioxidant rescue experiments.

### Contrast with a non-cardiac proteome under paraquat

Finally, we performed a secondary experimental end-point to further evaluate how the observed proteomic responses in the heart compare with the global response to paraquat in the animal. We collected the lungs from the cadavers of identical animals and subjected them also to isotope labeling tandem mass spectrometry analysis (**Figure S4a**). Both the lung and the heart are high energy-demand tissues that are susceptible to oxidative damage, and previous work has identified the lung to be among the tissues with highest accumulation of paraquat. We therefore examined whether the proteomic responses in the lung to paraquat treatment mimicked those in the heart.

At 10% FDR, we found a robust up-regulation of PDK4 (logFC: 3.4; P: 3.1e–4), CAT (logFC: 1.2; P: 2.3e–4), and HMOX1 (logFC: 5.7; P: 2.2e–4) in the lung, indicating paraquat also induced an acute oxidative stress response in the lung of the examined animals. Comparison of organ-specific responses is made challenging by the different proteome composition and dynamic range of concentration between the organs, and moreover, each tissue may be differentially sensitive to paraquat or reacts in different time scales. Nevertheless, we were surprised to find that the fold-changes of proteins commonly quantified in both the heart and the lungs following paraquat treatment showed little to no correlation to each other (*ρ* : 0.097), even when considering only proteins that were nominally differentially regulated in the heart (limma P ≤ 0.01) (*ρ* : 0.22). We observed a number of differentially regulated proteins in the lung that were not changed in the heart, including mitochondrial fission factor (MFF) (lung logFC: 2.03, P: 2.7e–4, adjusted P: 8.7e–2; heart logFC: 0.00, P: 0.99) and glypican 1 (GPC1) (lung logFC 4.5, P: 4.7e–8, adjusted P: 1.6e–4; heart logFC: –0.01, P: 0.99), which could form the basis for future investigations into cardiac-specific oxidative stress response elements.

## Summary

Protein differential expression analysis identified global changes in metabolism, antioxidant proteins, and proteostasis pathways that are consistent with a complex cardiac response to oxidative stress at the proteome level. A metabolic component of the mitochondrial unfolded protein response has been previously indicated in the literature and the data here suggest that in paraquat-injured hearts this is effectuated in part through the activation of PDK4. We also found PDK4 to be up-regulated in the lungs of the animals, but as this PDK isoform is not ubiquitously expressed, it would be interesting to assess the mechanism of mitochondrial response, if any, in other tissues. Lastly, we found a number of proteins with as-yet unclear function in the cardiovascular system that are acutely induced by oxidative stress and which merit further investigation.

With the experimental design incorporating subcellular fractionation, we found that paraquat preferentially induced changes in one fraction (S2) over the others in the heart. Differential detergent lysis is a common method to resolve partially overlapping protein fractions that correspond in part with specific cellular compartments [35]. It is now appreciated that different cellular compartments have distinct requirements and mechanisms for maintaining protein integrity [8,36]. The results here corroborate that paraquat primarily affects proteins in mitochondria and endoplasmic reticulum enriched compartments in the heart. Moreover, we found evidence that some proteins may redistribute in fractions upon paraquat challenge. These results therefore indicate that spatial dynamics may be a component of proteomic features during the acute stress response and echo the increasing number of reports that demonstrate proteome-wide spatial redistribution upon a variety stimuli [37].

Taken together, we report here a deep proteomics profiling dataset on the global protein expression changes of an acute oxidative stressor on the heart using isotope labelling tandem mass spectrometry. The results and data presented here may avail ongoing efforts to understand how the heart responds to oxidative stress and identify intervention targets.

## Methods

### Animal models and tissue collection

Twelve-week old male wild-type C57BL/6J mice were purchased from Jackson Laboratories (Bar Harbor, ME, USA) and housed in accordance with guidelines set by the National Institutes of Health (NIH) for the Care and Use of Laboratory Animals. The protocol for animal use was approved by the Animal Care and Use Committee of the University of Colorado School of Medicine. Animals were held in a temperature-controlled environment on a 12-hour light/dark cycle and supplied with pellet chow and water ad libitum. In the first animal experiment, mice were randomized into six treatment groups (n=4 per group each) and received either Dulbecco’s phosphate buffered saline (DPBS) (200 *µ*L, i.p.) or paraquat (N,N-dimethyl-4,4-bipyridinium dichloride) at three doses (10, 50, 75 mg/kg i.p.) and examined after 24 hours. One group was examined after 48 hours of 50 mg/kg paraquat i.p, and the last group received 2 mM Tempol (4-hydroxy-2,2,6,6-tetramethylpiperidin-1-oxyl) in drinking water for 24 hours ad libitum prior to paraquat (50 mg/kg i.p.) treatment and examined after 24 hours post-paraquat. Chemicals were purchased from Sigma unless specified.

In the second experiment, mice were randomized into four treatment groups (n=5 per group). Mice in vehicle and paraquat exposure groups received either DPBS (200 *µ*L, i.p.) or paraquat (50 mg/kg, i.p.) for 24 hours, respectively. Mice in the antioxidant treatment groups were administered with Tempol (2 mM in drinking water, ad libitum) or its triphenylphosphonium-conjugated lipophilic variant MitoTEMPO (0.7 mg/kg, i.p.) for 24 hours prior to administration with paraquat (50 mg/kg, i.p.) for 24 hours. Paraquat and MitoTEMPO solutions were prepared in DPBS. One group of paraquat-treated animals was euthanized prior to the experimental end-point; and the remaining four replicate groups were used in the experiments. Following exposures, mice were euthanized by CO_2_ displacement followed by cervical dislocation, and the heart and lungs were harvested. The harvested tissues were rinsed in PBS, snap-frozen immediately in liquid nitrogen, and stored at –80 ^°^C until analysis.

### Compartmental protein extraction and digestion

To extract proteins from subcellular fractions of the mouse tissues, harvested tissues were washed with PBS, blotted dry, and weighed, then homogenized using an Omni tissue homogenizer (Omni International, Kennesaw, GA, USA) using 3 pulses of 20 sec each at setting 4. Subcellular fractions were extracted using differential lysis and solubilization with a commercial Millipore Compartmental Protein Extraction Kit (Millipore) according to manufacturer’s instructions. Briefly, compartments S1, S2, and S3 were attained through detergents supplied as buffer C (KCl, glycerol, sodium orthovanadate), buffer N (NaCl, glycerol, sodium orthovanadate), and buffer M (KCl, sucrose, glycerol, sodium deoxycholate, NP-40, sodium orthovanadate). Following extraction, total protein concentration from each fraction was determined by bicinchoninic acid assay (Thermo Pierce). 150 *µ*g of protein was digested on a molecular weight filter. Briefly, samples were placed on 10 kDa MWCO polyethersulfone filters (Thermo Pierce), denatured with 8 M urea, and buffer exchanged into 100 mM triethylammonium bicarbonate (TEAB). Samples were reduced (55 ^°^C, 30 min) with 3 mM Tris(2-carboxyethyl)phosphine hydrochloride(TCEP-HCl) (Thermo Pierce) while shaking at 600 rpm in a thermomixer (Eppendorf). Samples were alkylated with 9 mM iodoacetamide (22 ^°^C, 30 min) in the dark with 600 rpm shaking, then digested on-filter with sequencing-grade trypsin (Promega) at a ratio of 50:1 (w/w) for 16 hr at 37 ^°^C, shaking at 600 rpm. Post-digestion peptide concentration was determined using a Pierce quantitative colorimetric peptide assay kit (Thermo Pierce) then labeled using tandem mass tag isobaric stable isotope labels (Thermo TMT10plex™; lot TA260585).

### Liquid chromatography and tandem mass spectrometry

We used a block multiplexing approach to design the randomization scheme and assign the comparison group samples to tandem mass tag channels. For each tissue, samples from all treatment groups and subcellular fractions were randomized to one of ten channels in one of six blocks using the RAND function in Excel with no control over randomization seed (**Figure S1**). Biological replicates within the same subcellular compartments were assigned to the same blocks where possible for batch correction. For accurate comparisons across blocks, two identical internal reference standards were also generated by combining 20 *µ*gof peptides pooled from nine representative samples within a labeling block, and randomized to two channels in each block. TMT10plex™ labeling for each block was performed according to manufacturer’s instructions. Briefly, TMT10plex™ reagents were reconstituted in 41 *µ*L of acetonitrile, combined with 10 *µ*g of peptide, and incubated for 1 hr at 22 ^°^C. The labeling reaction was quenched by incubation with 8 *µ*L of 5% hydroxylamine in 100mM TEAB for 30 min at 22 ^°^C, shaking at 600 rpm. The labeled peptides were subsequently combined, dried by vacuum centrifugation, and reconstituted in 300 *µ*L of 0.1% trifluoroacetic acid in LC-MS grade water. Samples were then subjected to offline reversed phase fractionation (8 fractions) using a Pierce High-pH Reversed Phase Fractionation Kit (Thermo Fisher) according to the manufacturer’s instructions for fractionation of TMT-labeled samples.

Liquid chromatography-tandem mass spectrometry was performed on fractionated labeled peptides. Second dimension liquid chromatography was performed online with an Easy-nLC 100 ultrahigh-pressure liquid chromatography (UPLC) nanoflow system (Thermo Fisher) connected via an EasySpray interface to a Thermo Q-Exactive HF mass spectrometer (Thermo Fisher) using an EasySpray PepMap C18 column (3 *µ*M particle size, 100 Å pore size, 75 *µ*m i.d. × 150 mm length; Thermo). 3 *µ*L of the peptide samples were injected via the autosampler. The nano-UPLC was run at 300 nL/min with the gradient of 0 to 105 min, 0 to 40% B, 105 to 110 min, 40 to 70% B, 110 to 115 min, 70 to 100% B, hold for 5 min, with solvent A being 0.1% v/v formic acid in water and solvent B being 80% v/v acetonitrile and 0.1% v/v formic acid. Typical MS1 survey scan was acquired at 60,000 resolving power in positive polarity in profile mode from 300 to 1650 m/z, lock mass, dynamic exclusion of 30 s, maximum injection time of 20 msec, and automatic gain control target of 3e6. MS2 scans were acquired on the top 15 ions with monoisotopic peak selection at 60,000 resolution, automatic gain control target of 2e5, maximum injection time of 100 ms, and isolation window of 1.4 m/z, with typical normalized collision-induced dissociation energy of 32.

The Thermo raw spectrum files were converted to the open source .mzML format [38] using ThermoRaw-FileParser v.1.2.0 [39] with the options (-g -f 2). MS2 spectra were searched using SEQUEST algorithm implemented in Comet (v.2019.01 rev.4) [40] against a .fasta database Uniprot/SwissProt *mus musculus* database accessed on 2020-03-19 (17,033 target entries) [41]. Comet search parameters followed conventional standards including: peptide_mass_tolerance=10, peptide_mass_units=2, isotope_error=3, num_enzyme_termini= 1, allowed_missed_cleavage=2, fragment_bin_tol = 0.02. Variable methionine oxidation (M +15.9949Da), static cysteine carbamidomethylation (C +57.021464 Da), and static tandem mass tag addition to lysine or the peptide N-terminus (nK +229.16293 Da) were specified. Peptides were reranked and assigned with confidence of identification using the Crux/Percolator v.3.2 [42] followed by protein inference, with peptides accepted for identification under 0.01 Percolator q-value.

### Protein quantification and statistical analysis

As the Crux/Percolator pipeline does not come with a native isobaric label quantification tool, we wrote a Python 3 tool, py-tmt-quant, to integrate the label channel intensities of each MS2 spectrum in the experiment. The software tool reads in the supplied Percolator result file and a directory of corresponding mzML files using the pymzml library, and returns an appended search result file with the channel intensities. It is compatible with MS2-level tandem mass tag experiments using TMT 0, 2, 6, 10, 11, and 16-plex tags and allows user-defined mass tolerance for integration (see Code Availability). Using py-tmt-quant we integrated the centroid peak intensity within 10 ppm of each channel m/z. For each peptide-charge combination, only the PSM with the best Percolator score per fraction is integrated. We corrected for isotope impurities using the contamination matrix from the manufacturer (**Figure S1B**). PSMs were filtered to exclude peptides from common contaminants (albumin, hemoglobins, and keratins), those without any TMT channel intensities, and those with no internal reference standard channel intensity, as well as those with low TMT channel intensities below 10% of total spectral intensity.

We performed sample-level dye correction through column sum normalization. We then performed block-wise correction using the ComBat batch correction function implemented in the sva package (v.3.35.2) [43] in R (v.3.6.3)/Bioconductor (v.3.10) [44]. Protein-level variances were normalized using the median values of the internal reference standard channel intensities across each block, and finally the effective total intensities were normalized using trimmed means of m-values with singleton pairing as implemented in the edgeR package (v.3.29.1) [45] in R/Bioconductor (**Figure S1C**). Statistics on differential protein expression were performed using the moderated t-test implemented in the limma package (v.3.43.5) correcting for duplicate correlations of subcellular fractions from identical animals [46], followed by the Benjamini-Hochberg procedure [47] to correct for multiple testing and calculate false discovery rates. Additional analysis on protein function was performed with the aid of StringDB [48], Pubpular [49], Reactome [50], and Enrichr [27].

## Data and Code Availability

Raw mass spectrometry files will be uploaded to ProteomeXchange with reviewer access credentials pending publication. The latest version of py-tmt-quant (v.0.3.0) will be accessible on Github.

## Disclosure

The authors declare no conflicts of interest.

## Acknowledgments

This study was supported in part by NIH NHLBI awards R00-HL127302, R01-HL141278 to M.L. and K99-HL144829 to E.L.; NRSA Postdoctoral Fellowship F32-HL149191 to Y.H.; the University of Colorado Postdoctoral Fellowship in Cardiovascular Research T32-HL007822 to V.D. and Y.H.; and the University of Colorado Consortium for Fibrosis Research and Translation Pilot Grant to M.L.

## Supplementary Figures and Tables

**Figure S1.**
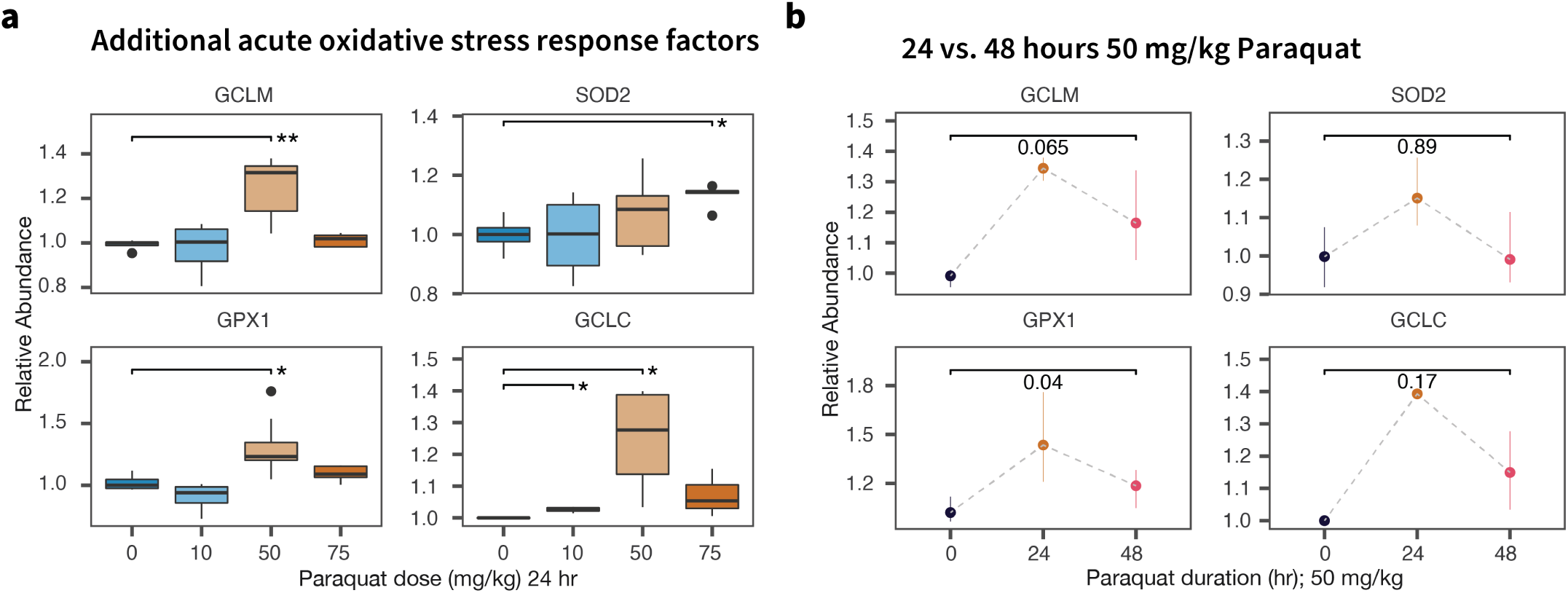
Paraquat induced changes of additional oxidative stress response proteins. **a**. Protein abundance of GCLM. GCLC, SOD2, and GPX1 at three doses of paraquat for 24 hours; *: t test P < 0.05;* * : 0.005. **b**. Comparison between 24 hr and 48 hr treatment of 50 mg/kg paraquat; Numbers: t test P values.

**Figure S2.**
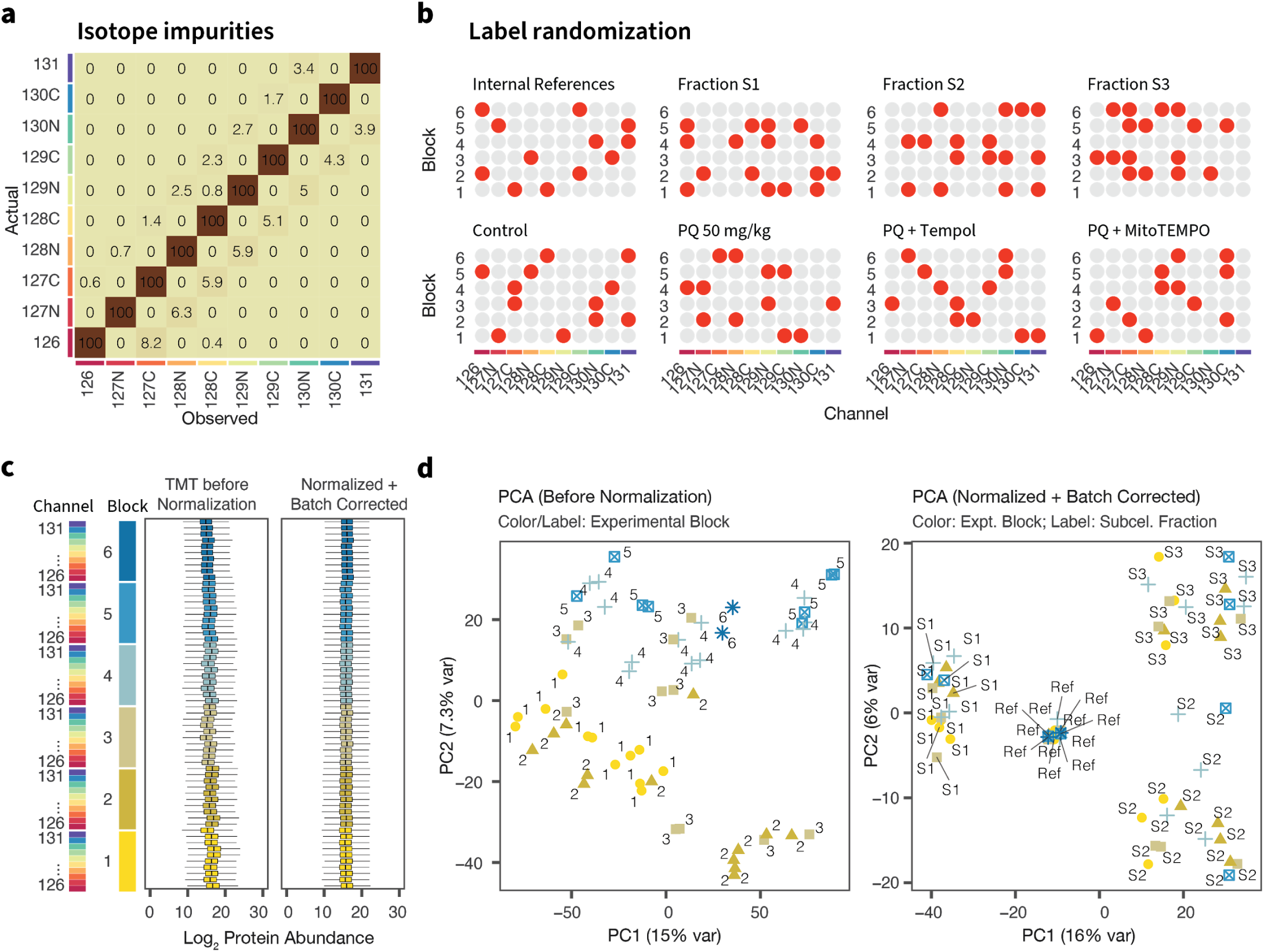
Isotope labeling tandem mass spectrometry analysis of normal and stressed hearts. **a**. Isotope impurity matrix for the tandem mass tag channels supplied by the manufacturer. **b**. Label assignment of samples in 6 experimental batches. Labels were randomized using a random number generator in Excel. **c**. Distribution of log label abundance before and after normalization and batch correction. **d**. Principal component analysis showing preferential grouping of channel intensities across experimental batches prior to normalization, and correction after.

**Figure S3.**
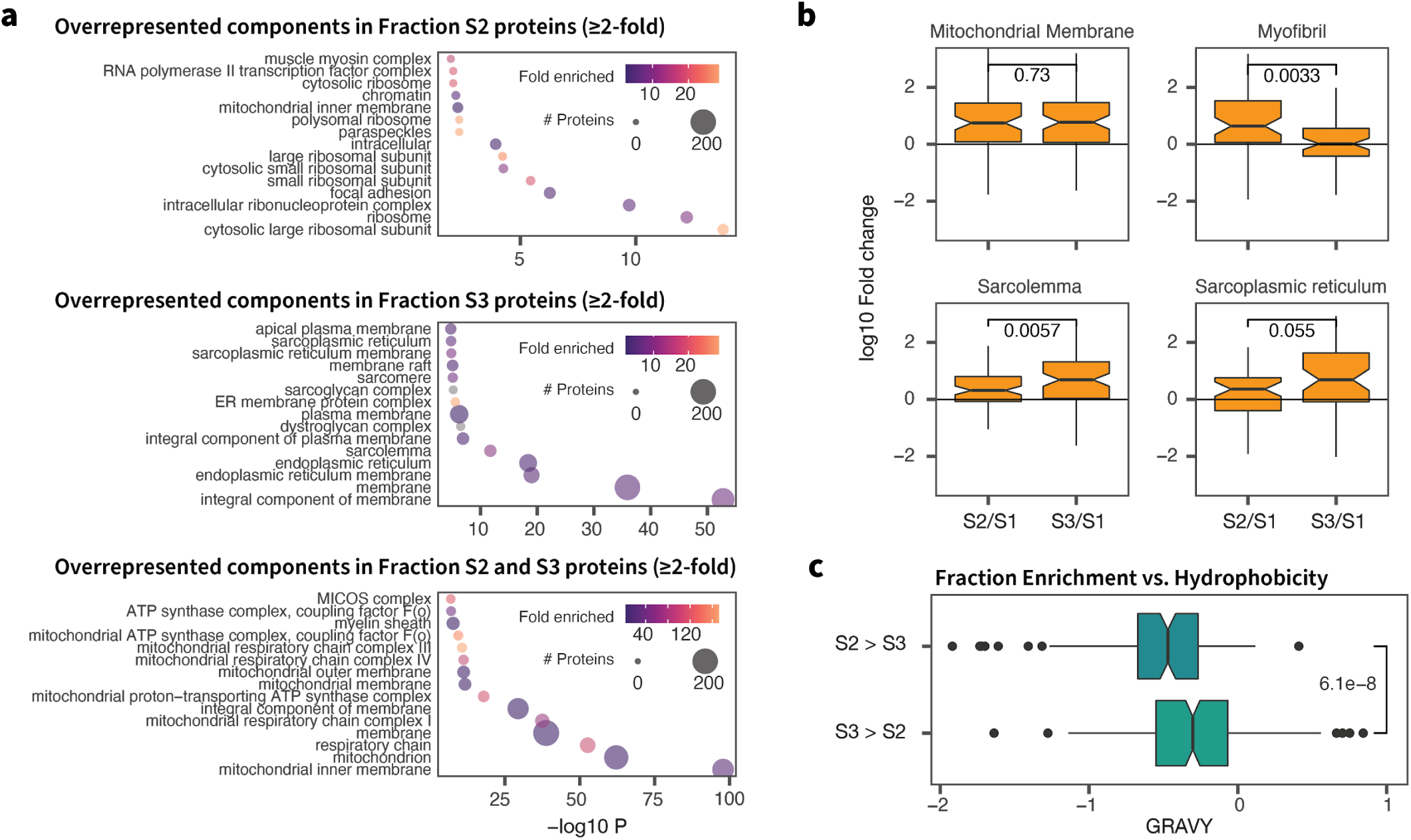
Properties of proteins enriched in subcellular fractions. **a**. Top 15 significantly enriched Gene Ontology cellular compartment terms in proteins in the S2 (top); S3 (middle); and both S2 and S3 fractions over S1. Data point size denotes number of proteins associated with a term; color denotes fold enrichment of term representation over background; x: –log P of hypergeometric test. **b**. Box plots showing relative enrichments in S2 and S3 fractions over S1 of proteins in the mitochondrial membrane, myofibril, sarcolemma, and sarcoplasmic reticulum. **c**. Grand average of hydrophobicity (GRAVY) scores of proteins that preferentially enrich in S2 vs. S3 fraction, indicating the S3 fraction contains more hydrophobic proteins.

**Figure S4.**
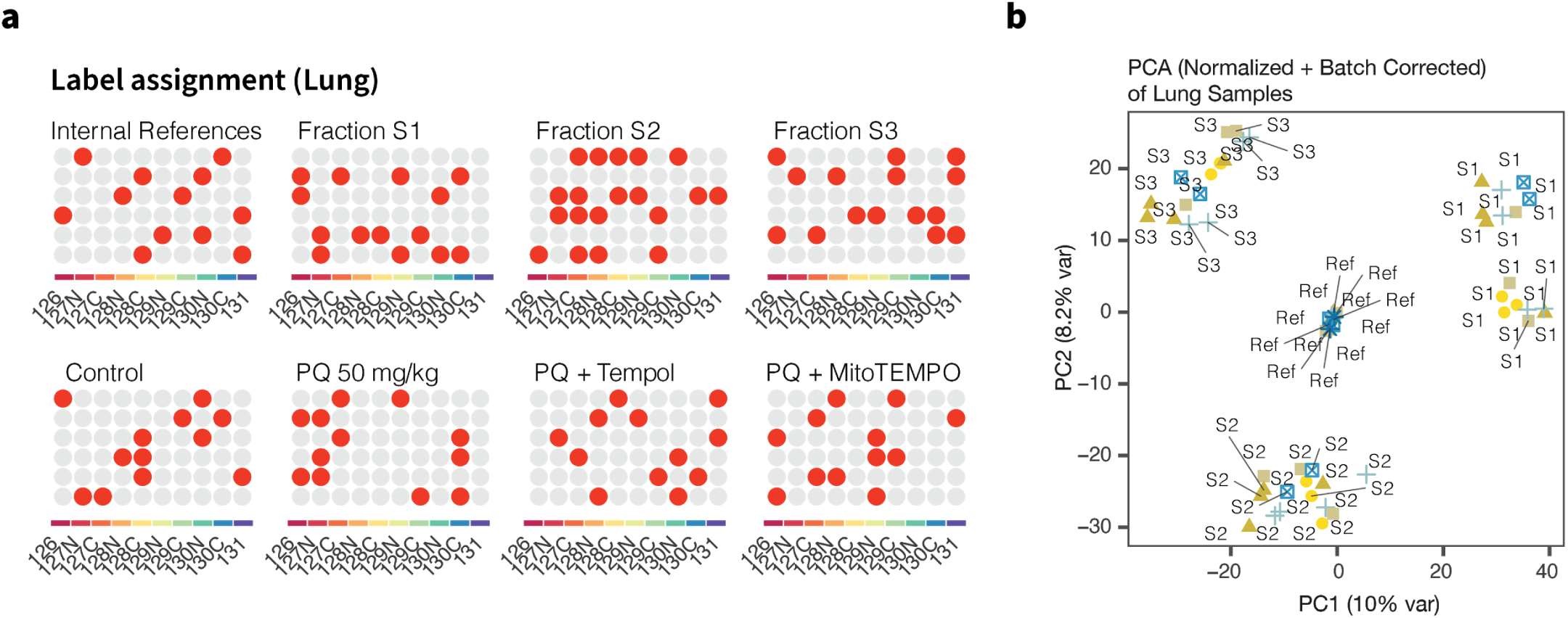
Isotope labeling tandem mass spectrometry analysis of normal and stressed lungs. **a**. Label assignment of samples across 6 experimental batches. Labels were randomized using a random number generator in Excel. **b**. Principal component analysis showing expected preferential grouping of samples by subcellular fractionation following normalization and batch correction.

## Notes

### Competing Interest Statement

The authors have declared no competing interest.

